# Enhanced human sensorimotor integration via self-modulation of the somatosensory activity

**DOI:** 10.1101/2024.10.20.619209

**Authors:** Seitaro Iwama, Takamasa Ueno, Tatsuro Fujimaki, Junichi Ushiba

## Abstract

Kinesthetic motor imagery, the mental rehearsal of movement and its associated sensations^1,2^, activates somatosensory and motor cortices^3,4^. Motor imagery improves performance in neurorehabilitation^5^, sports^6^, and instrument play^7^. However, the mechanism that enhances performance through repetitive motor imagery remains unclear. Here, we combined individually tailored motor imagery training combined with closed-loop neurofeedback training, neurophysiology, and behavioral assessment to characterize how the training can modulate the somatosensory system and improve performance. The closed-loop training using real-time feedback of human electroencephalogram (EEG) signals enhanced participants’ self-modulation ability of intrinsic neural oscillations in the primary somatosensory cortex (S1) within 30 minutes. Further, the short-term reorganization in S1 was corroborated by the post-training reactivity enhancement revealed by changes in somatosensory evoked potential (SEP) amplitude of early component originating from S1^8^. Meanwhile those derived from peripheral sensory fibers were maintained, suggesting that the closed-loop training manipulated cortical activities. Behavioral evaluation demonstrated improved performance during keyboard touch-typing indexed by resolved the speed-accuracy trade-off^9^. Collectively, our results provide evidence that mentally rehearsed movement during kinesthetic motor imagery induces functional reorganization of S1 activities that can contribute to performance improvement.

## Results

In the present study, we hypothesized that the repetitive kinesthetic motor imagery combined with neurofeedback training would be beneficial for the subsequent motor performance because it rehearses task-related primary somatosensory and motor cortex (S1 and M1) activation and leads to functional remodeling of the somatosensory pathways. To this end, we asked if induced sensorimotor plasticity through the short-term motor imagery training is boosted by the closed-loop feedback, manipulates S1 reactivity, which is tested by the somatosensory evoked potential (SEP), and improves sensorimotor performance in healthy humans.

To train for the self-modulation of sensorimotor neural excitability, we employed individually tailored closed-loop training. During the training, we provided virtual finger movement based on real-time estimated the sensorimotor rhythm (SMR) amplitude^10^ (Figure 1a). If the self-modulated SMR can manipulate the somatosensory pathways, the SEP amplitude change would accompany SMR modulation.

**Figure 1.**
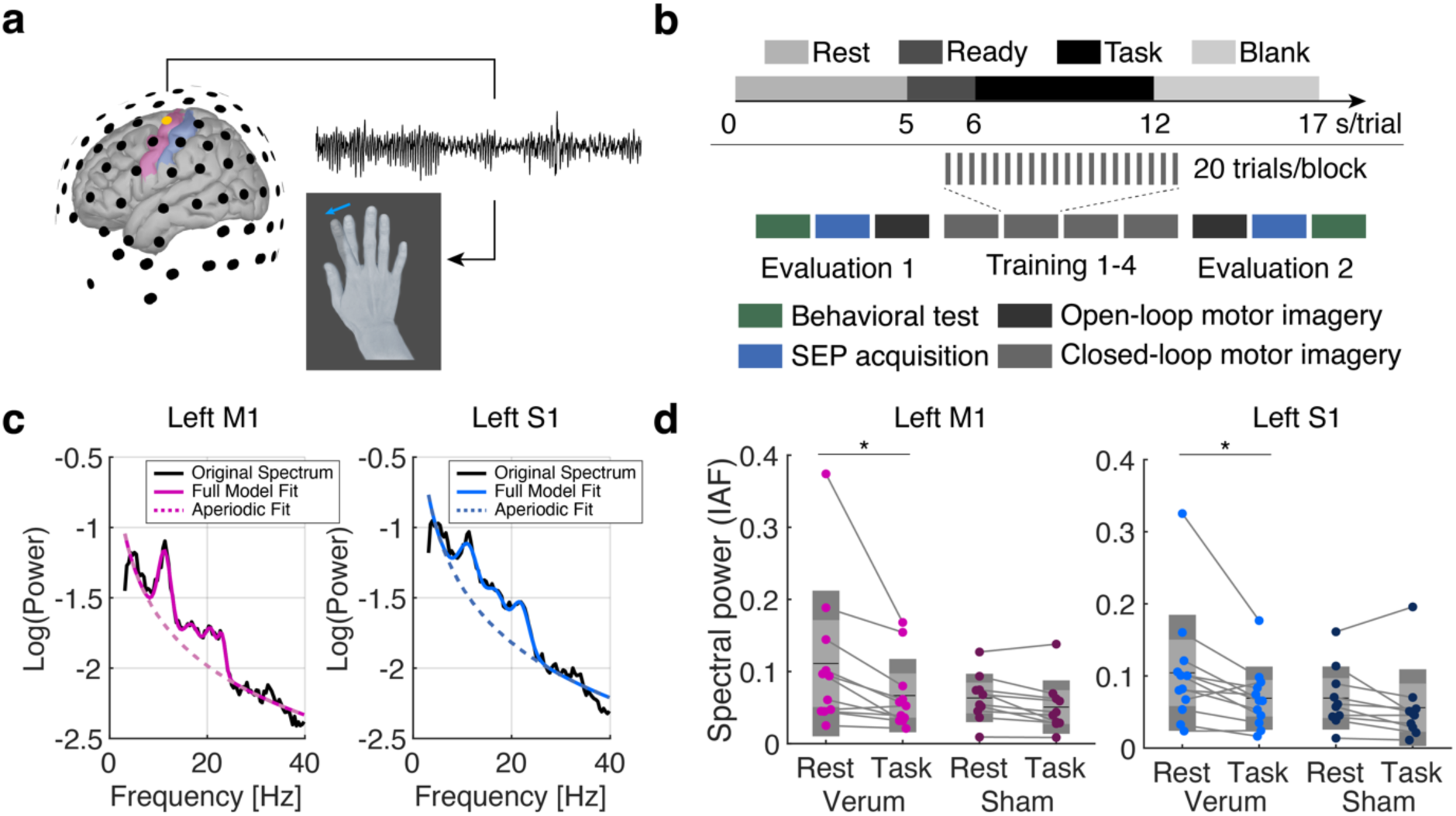
Closed-loop sensorimotor activity feedback experiment. **a** Schematic of experiment. Real-time processed scalp electroencephalogram (EEG) signals derived from the left sensorimotor cortex were associated with the movement of right index finger abduction shown on the screen. The range of motion represents EEG spectral power attenuation during motor imagery task. **b** Experiment protocol. The upper panel indicates time course of a motor imagery trial which comprised of four periods: Rest, Ready, Task, and Blank. The entire trial lasted 17 seconds and 20 trials were conducted in a block. The experimental procedure includes two evaluation (open-loop) and four training (closed-loop) blocks. These blocks were flanked with behavioral test, and somatosensory evoked potential acquisition. **c** Power spectral density (PSD) plots for the left primary motor and sensory cortices (M1 and S1). The black solid lines indicate the original spectrum, and the colored indicate the full model fit acquired by the *specparam* algorithm ^13^. The dotted lines represent the aperiodic fit. **d** Spectral power peak at individual alpha frequency (IAF) for the left M1 and S1.

### Training-induced neural activity modulation was frequency and region specific

We recruited 39 participants for the main experiment and data from 22 participants were subjected to main analysis. We excluded relatively larger sample size due to the high drop rate for the aversion to the median nerve stimulation and compromised quality for the SEP data (See Material and Methods for the criteria).

The participants were asked to perform motor imagery of index finger abduction and instructed to kinesthetically imagine contracting the right first dorsal interosseous muscle (FDI muscle) during which the joint angle of the index finger displayed on a monitor were controlled based on the spectral power of EEG signals at individually adjusted alpha frequency (IAF), derived from the contralateral sensorimotor cortex (i.e., C3, Figure 1a). Namely, the finger was configured to show abduction when the IAF spectral power at the C3 channel was attenuated compared to a 5-second resting state period before a 6-second motor imagery period (Figure 1b).

The participants were allocated to one of two groups: the verum group, where they observed the images of finger movement linked with their own scalp EEG data, and the Sham group, where they observed those linked with previous data from others (Verum: *N* = 12; Sham: *N* = 10). The group allocation was randomized and single-blinded throughout the study. No participants reported adverse effects throughout and after the experiment.

As a first step of the offline analysis, we estimated the cortical source using the high-density EEG signals and differently assessed the M1 and S1 activities^11,12^. After calculating the power spectral density derived from the contralateral M1 and S1, IAF was identified using the spectral parameterization algorithm (*Specparam*^13^, Figure 1c). The spectral power at the IAF was compared with a mixed repeated-measures ANOVA (rmANOVA) with condition (Rest or Task), and group (Verum or Sham) effects (Figure 1d). We found a significant main effect of condition in both left M1 and S1 data (M1: *F* = 8.94, *p* = 0.008, *η*² = 0.051; S1: *F* = 9.62, *p* = 0.006, *η*² = 0.045). Post-hoc *t*-tests revealed a significant difference between the rest and task periods during training blocks in the two areas in the verum group (Paired *t*-test with Bonferroni correction, M1: *t* = 3.37, corrected-*p* = 0.019, *d* = 0.72; S1: *t* = 3.34, corrected-*p* = 0.02, *d* = 0.62). Since attenuated spectral power around SM1 reflects desynchronized rhythmic activities due to cortical excitability change^4,14^, the task-related power attenuation indicates the self-modulation of SM1 activity was successfully induced during motor imagery training^5,15^. A significant main effect of group was also found in the beta-band power, but not evident in the theta band (Supplementary Figure 1ab). In addition, the between-group difference in IAF, and the broadband aperiodic component were not evident, suggesting that the oscillatory spectral power related to the SMR component used in the real-time feedback was modulated during the closed-loop training.

To confirm whether the periodic component of the power spectral density (PSD) was modulated during the training, the spectral peak height at IAF was compared. A significant group-difference during the rest period in training blocks was found (Supplementary Figure 2a, Two-sample *t*-test, M1: *t* = 2.79, *p* = 0.0014, *d* = 1.15; S1: *t* = 2.74, *p* = 0.0015, *d* = 1.14) while no other frequency bands exhibited systematic between-group difference (Supplementary Figure 2bc). Collectively, the periodic power change indicates that closed-loop SMR control manipulates neural population oscillating at the targeted frequency.

### Modulation depth of the sensorimotor oscillatory activities was augmented by the closed-loop prosthetic control training

Given that the verum group indicated successful SMR modulation during training, we evaluated the endogenous modulation depth of SMR during the motor imagery task in the open-loop motor imagery blocks (Figure 2). To this end, we quantified event-related spectral perturbation (ERSP) magnitude at IAF for each participant, which represents the within-trial normalized magnitude of spectral power modulation calculated Rest period data as baseline. A significant decrease in the ERSP magnitude around 8-30 Hz was found in both left M1 and S1, indicating that motor imagery tasks induce significant self-modulated SMR in the Verum group (Figure 2a, One-sample *t*-test with Bonferroni correction). The significant areas of between-group comparisons for SMR were localized around SM1 in the left hemisphere (Fig 2b, Two-sample *t*-test with cluster-level false-discovery correction). Meanwhile, the sham group did not exhibit systematic modulation across participants. Collectively, participants in the verum group performed kinesthetic motor imagery accompanied with the significant activity modulation in SM1, despite its difficulty for the naïve participants^4,14,15^.

**Figure 2.**
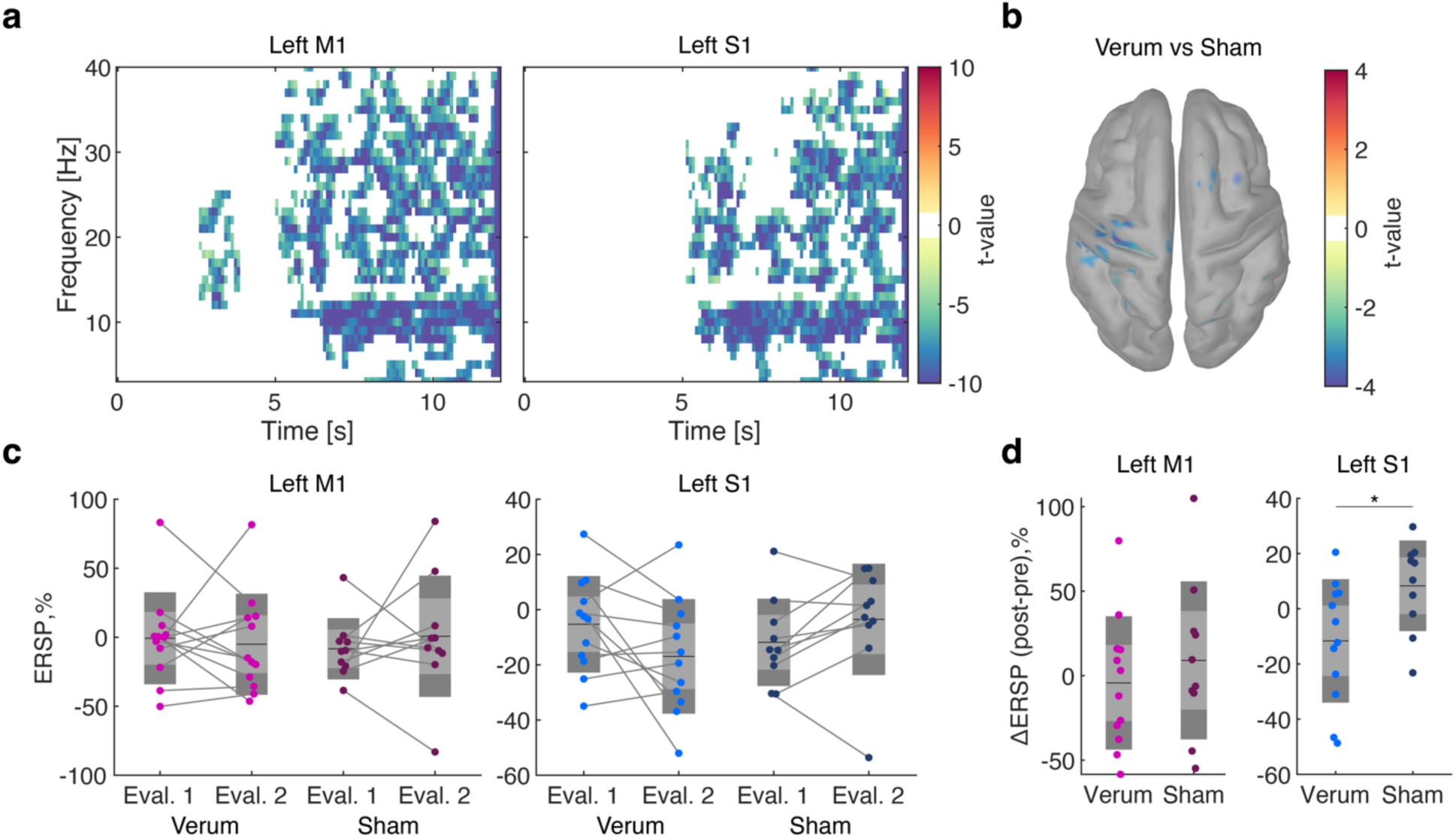
Time-frequency representations and comparison of event-related spectral perturbations (ERSP) between verum and sham groups. **a.** Time-frequency representation of power changes in the left M1 and S1 during the motor imagery task. The clusters exceeding the significance threshold (*p* < 0.0001) were colored with their t-values. Negative *t*-values represent significant spectral power attenuation compared to the rest period. **b.** Spatial distributions of significant *t*-values indicating difference between verum and Sham groups. **c.** Comparison of event-related spectral perturbation (ERSP) during Evaluation 1 and Evaluation 2 for verum and Sham groups in the targeted regions. Each dot represents an individual participant, and lines connect the same participant’s data across evaluations. The dark grey areas, light grey areas, and the black line represent 1 SD, 95% confidence interval, and mean values, respectively. **d.** Changes in ERSP from Evaluation 1 to 2 for the verum and sham groups in the left M1 and S1. A significantly prominent spectral power attenuation was found in the left S1 in the verum group.

Next, to compare the task-related modulation depth of SMR magnitude, the block-average ERSP magnitudes were subjected to a mixed rmANOVA (Figure 2c). The rmANOVA revealed a significant interaction of group and time effects in the S1 ERSP magnitude in the trained hemisphere (rmANOVA, M1: *F* = 0.54, *p* = 0.47; S1: *F* = 5.61, *p* = 0.028, *η*² = 0.07). Post-hoc two-sample *t*-tests for the changes in ERSP magnitude from Evaluation 1 to 2 revealed that the verum group exhibited a significant decrease in ERSP magnitude compared to the sham group (Figure 2d, Two-sample *t*-test, M1: *t* = -0.72, *p* = 0.48; S1: *t* = -2.44, *p* = 0.024, *d* = 1.03). This suggests that short-term kinesthetic motor imagery induced enhanced S1 activities without any somatosensory feedback, consistent with the recent findings in the non-invasive functional neuroimaging data during motor preparation^16,17^ as well as the single neuronal activity derived from S1^18,19^.

### Neurofeedback-induced changes in the somatosensory evoked response

To further investigate the somatosensory reactivity manipulated by the closed-loop motor imagery intervention, we assessed the changes in SEP magnitude derived from multiple locations (Figure 3a). The ascending somatosensory signals evoked by right median nerve stimulation were visualized as N9, N13 and N20 components, respectively. Each of the components reflect difference neuronal reactivity^20,21^: N9 represents the brachial plexus response, N13 represents the cervical spinal cord response, and N20 represents the early component of the S1 response, specifically Brodmann’s 3b area^8,21^.

**Figure 3.**
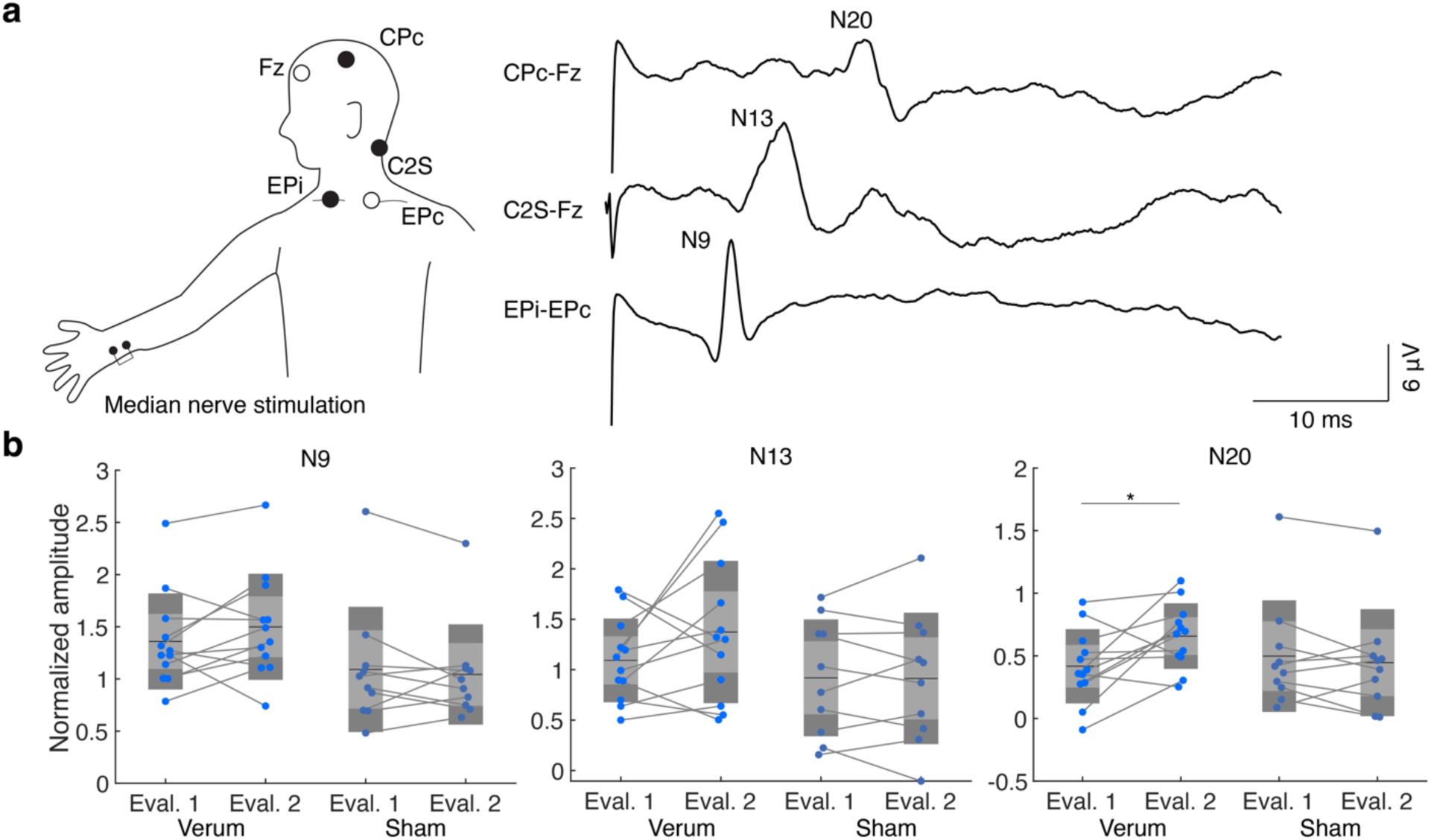
Somatosensory evoked potentials (SEP) results. **a.** Schematic representation of electrode placement for recording SEPs elicited by right median nerve stimulation. Electrode positions were Fz, CPc, C2S, EPi, and EPc. The SEP waveforms show typical peaks (N20, N13, and N9) recorded at different electrode montage (CPc-Fz, C2S-Fz, and EPi-EPc). **b** Normalized amplitudes of SEP components (N9, N13, N20) during Evaluation 1 and Evaluation 2 for the verum and sham groups. Each dot represents an individual participant, and lines connect the same participant’s data across evaluations. A significant interaction of time and group effect was found in the N 20 component, and post-hoc *t*-test with Bonferroni correction revealed a significant increase in amplitude in the verum group (rmANOVA: N9: *F* = 2.18, *p* = 0.16; N13: *F* = 1.75, *p* = 0.20; N20: *F* = 6.49, *p* = 0.019, *η²* = 0.04; post-hoc *t*-test for N20 data: Paired *t*-test with Bonferroni correction, Verum: *t* = -3.10, corrected-*p* = 0.033, *d* = -0.68, Sham: *t =* 0.614, corrected-*p* = 1.0).

To identify changes in SEP amplitude induced by the training, three components were subjected to the rmANOVA (Figure3b). We found a significant interaction of time and group in the N20 component (rmANOVA, N9: *F* = 2.18, *p* = 0.16; N13: *F* = 1.75, *p* = 0.20; N20: *F* = 6.49, *p* = 0.019, *η*² = 0.04). Post-hoc *t*-tests revealed a significant increase in the amplitude in the Verum group (Paired *t*-test with Bonferroni correction, *t* = -3.10, *p* = 0.033, *d* = -0.68). Meanwhile other components did not exhibit significant interaction or main effects (All statistical results shown in Supplementary Information). Given that N20 originate from S1, S1 reactivity was significantly manipulated by the closed-loop motor imagery training without any somatosensory feedback.

### Improved sensorimotor task performance through motor imagery practice

The closed-loop motor imagery training without any somatosensory feedback manipulated both oscillatory S1 activity modulation during motor imagery in open-loop condition, and S1 reactivity to the somatosensory stimuli. If the motor imagery training can act as the rehearsal of S1 activation during motor tasks, the training could augment sensorimotor processing in which both M1 and S1 are actively involved.

To corroborate the hypothesis, we tested the behavioral performance during a motor task requiring sensorimotor integration process. Specifically, we employed a keyboard touch-typing task^22,24^ to quantify post-training meditation of speed-accuracy trade-off. Since the somatosensory function can be used to execute rapid and accurate finger movement, we calibrated the task difficulty to adjust the individual typing performance. We found no evidence in the significant difference group-level task difficulty, measured by the rank and number of letters of selected words (Two-sample *t*-test, word rank: *t* = 1.30, *p* = 0.20, log *BF*_10_ = −0.43; word letters: *t* = 1.94, *p* = 0.06, log *BF*_10_ = 0.32). Participants underwent 5 blocks of the typing task, with 30 words presented in each block, using a keyboard without key printing.

A trial began with the time limit presentation, the three conditions (2, 4 and 6 seconds) were presented in pseudo-randomized order (Figure 4a). The word presentation period lasted for the time limit and the blank period followed. A trial was counted as success when participants typed the given word without an error in each time limit. We calculated the accuracy for each condition in Evaluation 1 and 2 (Figure 4b). Then, the speed-accuracy trade-off curve was estimated by fitting the following formula, *y* = *a exp*(−*bx*) + *c* where *x* and *y* represent the accuracy and condition, respectively. We fitted parameters *a, b, and c* and the saturation speed *b* was subjected to a mixed rmANOVA with time and group effects (Figure 4c). A significant interaction of time and group effects were found (*F* = 6.06, *p* = 0.023, *η*² = 0.096). The post-hoc t-test revealed a significant increase in the saturation speed in the verum group after training (Paired *t*-test with Bonferroni correction, *t* = -3.31, corrected-*p* = 0.02, *d* = -1.21), suggesting that the sensorimotor processing was modulated through the closed-loop motor imagery training. Meanwhile we did not find a systematic change in other fitted parameters (Supplementary Figure 3).

**Figure 4:**
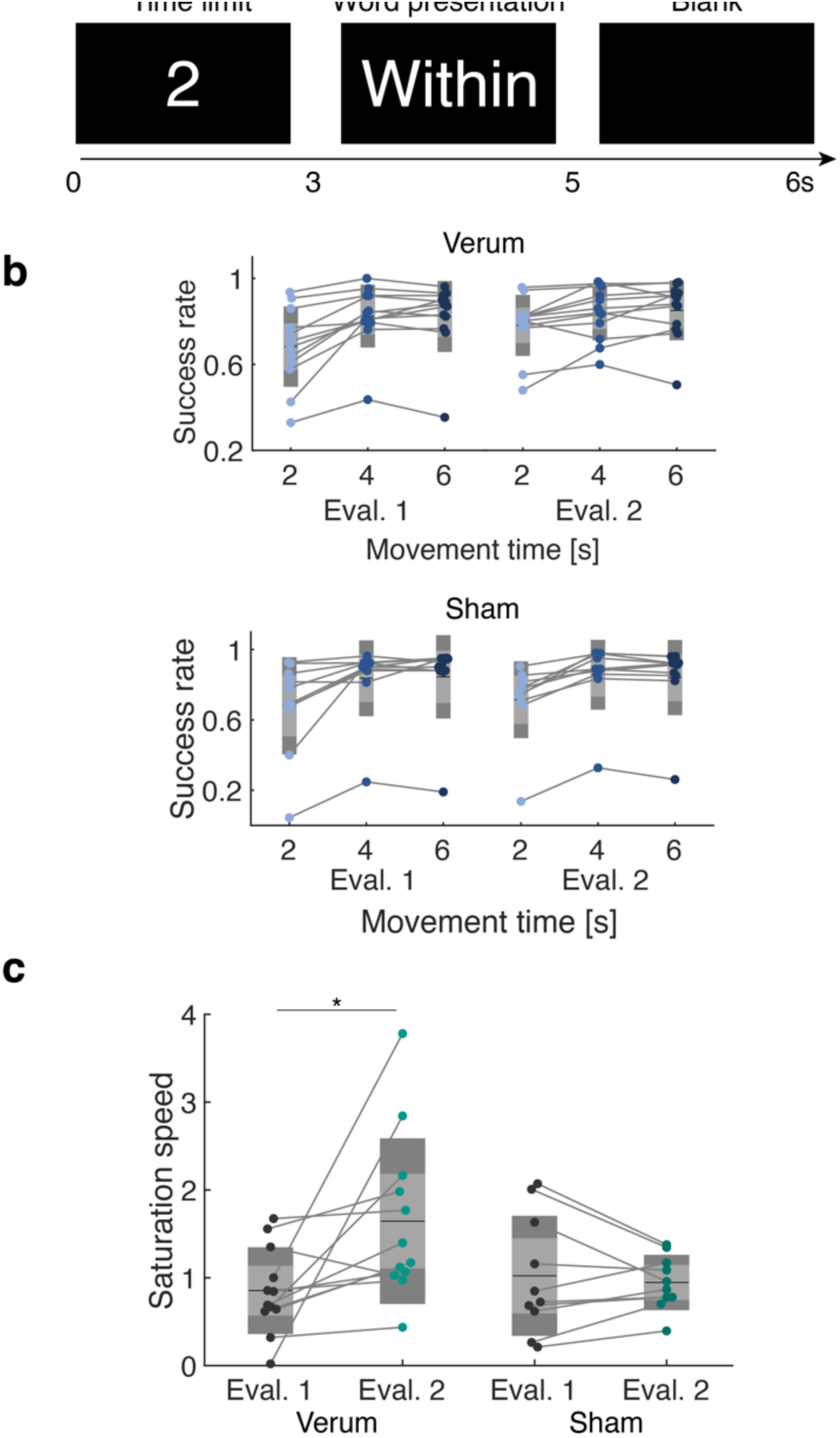
Behavioral test results. **a.** Schematic representation of a single trial in the experiment. The trial includes three periods: a 2-second time limit display, word presentation. and a 1-second blank. The duration of word presentation finished when participants failed to type the correct key. **b.** Success rates of participants in the verum and sham groups during Evaluation 1 and Evaluation 2 at different movement times (2, 4, and 6 seconds). Each dot represents an individual participant, and lines connect the same participant’s data across evaluations. **c.** Comparison of saturation speed (the speed at which the success rate plateaus) between Evaluation 1 and Evaluation 2 for the verum and sham groups. A mixed rmANOVA revealed significant interaction of time and group effects. The post-hoc *t*-test with Bonferroni correction revealed a significant increase in the Verum group (rmANOVA: *F* = 6.06, *p* =0.023, *η*² = 0.096; *t*-test: *t* = -3.31, *p* = 0.02, d = -1.21).

## Discussion

Human beings are adept at imagining the consequences of their actions and refining their motor plans to achieve manual dexterity. This skill perhaps leads to a rich repertoire of motor patterns and, ultimately, to human prosperity. To this end, humans employ kinesthetic motor imagery to engage the neural repertoires involved in a motor task without execution. Regardless of the absence of somatosensation, neuroimaging data including invasive electrophysiological recordings indicate task-related activation of the SM1 during motor imagery, putatively implicated in simulating the representation during movement initiation and preparation^17,23,25,26^. However, the direct link between the somatosensory activation during the kinesthetic motor imagery and subsequent performance improvement was unclear.

In the present study, we demonstrated that closed-loop motor imagery training specifically modulated the targeted sensorimotor cortex and significantly improved behavioral performance during a touch-typing task as indicated by the speed-accuracy trade-off. Furthermore, the skill acquisition of sensorimotor activity self-modulation resulted in a significant change in the N20 component of SEPs, indicating enhanced sensory reactivity in the S1^27–29^. These findings suggest enhanced somatosensory function, on which successful motor control is highly dependent^29^. We also found that the training effects in part generalized to the open-loop blocks during evaluation, where no feedback was provided. It indicates that the self-modulation ability of population-level S1 activity trained through the closed-loop motor imagery was not confined to the feedback blocks alone but also extended to the open-loop condition, and the training maintained significant SMR modulation depth. This suggests that the training induced lasting changes in sensorimotor cortex excitability that persisted beyond the immediate training context^30,31^. These observations are consistent with reports that manipulating M1 activity using a closed-loop neurofeedback paradigm influences corticospinal excitability^10,30,32,33^, indicating the potential utility of closed-loop motor imagery to enhance sensorimotor activities before the actual sensorimotor behavior is required^34^.

Our study has three limitations: the single-blind study design, the absence of an open-loop (i.e., no feedback) group, and the limited duration of the training period. First, the single-blind design may introduce bias, as experimenters were aware of their group assignments. While this design might influence subjective judgments, it is unlikely to affect the objective neurophysiological measures such as SEPs and SMR, which were the primary outcomes of this study. Additionally, the randomization of participants into groups helps mitigate potential biases and they were not informed about the existence of the other group. Therefore, it is not likely that the placebo and nocebo effects lead the result. Second, the absence of a purely open-loop group means it is unable to entirely separate the effects of motor imagery from those of feedback. Despite this, the significant differences observed between the verum and sham groups provide strong evidence that the feedback component plays a crucial role in modulating sensorimotor rhythms. Future studies incorporating a no-feedback group would clarify the specific contributions of feedback. Last, the short duration of the training period limits our understanding of long-term effects and consolidation of sensorimotor skills. However, the robust changes observed within this brief period suggest that even short-term interventions can induce apparent neural plasticity. Extending the duration of the training in future research will help determine the sustainability of these effects and their potential for long-term rehabilitation outcomes.

## Materials and Methods

### Participants

We recruited 39 participants for the main experiment and data from 22 participants were subjected to main analysis (17 males and 5 females; Age: 23.62 ± 3.4, All right-handed). We excluded relatively larger sample size due to the high drop rate for the aversion to the median nerve stimulation. Written informed consent to participate in the present study was obtained from every participant.

### Experiment procedure

All experiment protocol was approved by the Ethics Committee of the Faculty of Science and Technology, Keio University (IRB approval number: 2023-128) and performed in accordance with the Declaration of Helsinki. No adverse effects were reported throughout study.

The experiment comprised three phases: pre-training evaluation (Evaluation 1), training and post-training evaluation (Evaluation 2) as shown in Figure 1b. In the evaluation phase, participants underwent behavioral test, SEP acquisition, and the open-loop motor imagery block. In the training phase, participants experienced four blocks of closed-loop motor imagery training.

### Behavioral test

As a keyboard typing task, we used the English words listed in the General Service List^35^ (GSL), ensuring that non-native English speakers could recognize the words within the movement time limit. Participants performed a typing speed assessment before Evaluation 1 using a web service to quantify their characters per minute (CPM, https://typing-speed-test.aoeu.eu/). The CPM was used to determine the maximum word length. We used words with 80-100% of the characters that each participant was capable of typing within the given movement time for each condition.

### Somatosensory evoked potential acquisition

The SEP data were acquired to evaluate the reactivity of S1 by stimulating the right median nerve using Neuropack X1 (Nihon Kohden, Tokyo, Japan). The stimulation trigger and analog data derived from electrodes on CPc, Fz, C2S, EPi and EPc were sampled using the setup and recorded using an analog-digital converter (USB-6259, National Instruments, Austin, U.S.A.) at the sampling rate of 10000 Hz. The acquired signal data were filtered using a highpass filter (3 Hz) and a lowpass filter (500 Hz)^20^.

The stimulation intensity was adjusted to the individual motor threshold (MT). To determine MT, the sensory threshold was firstly identified by increasing the intensity until participants were aware of stimulus presentation, Then, MT was identified at the minimum intensity which elicit stimulation-evoked thumb movement. The MT determined in the Evaluation 1 was also used in Evaluation 2 to avoid confounding the sensory plasticity with stimulation intensity changes. In each Evaluation block, 1000 stimulations were presented at 2 Hz frequency.

### Kinesthetic motor imagery task

In the motor imagery blocks, we asked participants to perform kinesthetic motor imagery, that is to mentally rehearse body movement^2^. The participants performed the imagery of right index finger abduction. At the beginning of the experiment, participants were verbally instructed how to perform the kinesthetic motor imagery. The content of verbal instruction was consistent across participants using a script (See Supplementary Information).

In the open-and closed-loop motor imagery blocks, participants performed 20 trials in which a 5-second rest and a 1-second ready period were followed by a 6-second task period. During the task periods, participants were asked to perform the kinesthetic motor imagery as instructed. The absence of overt movement during the training was confirmed by the visual inspection.

### Real-time sensorimotor rhythm amplitude neurofeedback

In the closed-loop blocks, participants received the neurofeedback of SMR amplitude in real-time through virtual right index finger movement shown on the display in front of participants^15,31,34^. The spectral power of SMR was real-time calculated and associated with the Metacarpo Phalangeal joint angle (Figure 1a). At the beginning of the experiment, we captured the abduction of the right index finger using a video camera and extracted 20 frames from the movie data. The frame of the more abducted finger was presented if spectral power attenuation of SMR was found in the electrode over the left SM1.

To determine the frame to present, we analyzed the online acquired scalp EEG data. The experiment setup was employed in the previous EEG-based neurofeedback studies^15,31,36^. During the motor imagery blocks, we used the Electrical Geodesics system with 128-channel HydroCel Geodesic Sensor Net (GES 400, Electrical Geodesics, Inc.). EEG electrode impedance levels were kept below 50 kΩ throughout the experiment. A 100 Hz online low pass filter was applied to recorded data at a sampling rate of 1000 Hz. The EEG signals were subjected to online analysis to calculate SMR magnitude around contralateral SM1. The EEG signals were spatially filtered using a large Laplacian filter centered on C3 in the extended 10-20 system^37,38^. Every 100 ms, EEG signals of the latest 5 s were subjected to a third-order Butterworth bandpass (3-70 Hz) and notch (50 Hz) filter. After forward and backward filtering, the latest 1s data were subjected to Fourier transform with a Hanning window function to calculate signal strength in frequency space. This short-term Fourier transform analysis (STFT) resulted in a spectral band power. Using the signal power derived from the C3 electrode, SMR spectral power modulation was quantified as the size of ERSP using the following formula^14^:

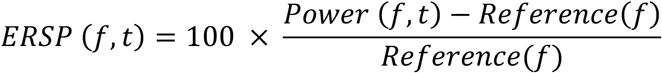

where *f* is frequency, *t* is time, *Power* is the signal strength, and *Reference* is the averaged power at the rest period. Then, SMR changes at the alpha-band (8-13 Hz) calculated by averaging the ERSP magnitude. The frequency bands used for online feedback were calibrated for each participant using frequency of interest (FOI) within the alpha-band. The FOI was determined from averaged ERSP during the open-loop block in Evaluation 1. Specifically, frequency that exhibited the most prominent negative at Task period in the alpha band was set as frequency of interest^15,31,39^. The online calculated SMR magnitude at FOI was averaged for the latest 10 samples to determine instantaneous cortical activity without being affected by signal flicker^15,31,34^.

The online calculated SMR magnitude at FOI was used for the neurofeedback. The 25th to 75th percentiles of SMR in the open-loop block in Evaluation 1 were used to map the given SMR magnitude and visual feedback. The number of frame where stronger SMR magnitude (25th percentile) was associated with the abducted index finger (the 20th frame).

In the sham group, participants received yoked-sham neurofeedback where the provided feedback visual stimuli were associated with the SMR magnitude calculated for the previously measured EEG signal derived from the other participant^40^. The identical procedure was conducted to determine the frame of the movie to present. Post-experiment debriefing did not reveal any participants who was aware of the intervention characteristic.

### Offline EEG spectral power analysis

In the offline analysis on the whole-head scalp EEG data, we computed the cortical source activities using sLORETA algorithm^12^ implemented in Brainstorm toolbox^11^. The estimated activities within M1 and S1 were subjected to the bandpass filtering and STFT analysis identical to the online processing.

To parameterize the power spectra, we used specparam algorithm^13^ and acquired IAF profiles and the aperiodic exponent which reflects the broadband 1/f component in the PSD. The ERSP values at IAF were computed for the offline analysis. In addition, whole-brain distributions of spectral power change were separately computed for each participant and averaged across the training blocks.

### Offline SEP analysis

To calculate the amplitude of each SEP component, the acquired time series data were epoched into 600 ms data where each stimulus onset was centered. Then, the averaged data across epochs were z-scored using the average and standard deviation of amplitude before stimulus. Then, N9, N13 and N20 components were searched by finding the extrema within the range of ±3 ms of each latency.

### Behavioral performance analysis

For each block, a successful trial was counted if the presented word was entered without error within the time limit, and the success rate for each movement time was calculated. Then, curve fitting using nonlinear least square algorithm was conducted for each participant success rate data using the following formula, *y* = *a exp*(−*bx*) + *c* where *y* and *x* represent the accuracy and condition, respectively. We fitted parameters *a*, *b*, *a,b, and c* and the parameter *b* was used to quantify the saturation speed^41^.

### Statistical analysis

Statistical tests were conducted using JASP^42^. EEG-SMR features, SEP amplitude, and behavioral performance were analyzed using a mixed rmANOVA with Time and Group factors. The sphericity was checked with the Mauchly’s test of sphericity for rmANOVA with more than two levels per factor. The post-hoc analysis was conducted using *t*-test with Bonferroni correction. To ensure the absence of strong evidence in the alternative hypothesis, we additionally calculated Bayes factor of the null- and alternative hypothesis.

## Acknowledgements

We thank Dr. F. Iwane for feedback on this paper. This study was supported by JST, PRESTO Grant Number JPMJPR23I1, Japan and JSPS KAKENHI Grant Number 20H05923, Japan. The authors thank Shoko Tonomoto Aya Kamiya and Sayoko Ishii for their assistance.

## Author contributions

Conceptualization, S.I., and J.U.; Methodology, S.I., and T.U.; Software, S.I., T.U. and T.F.; Validation, S.I.; Formal Analysis, S.I.; Investigation, S.I., T.U. and J.U. Resources, S.I. and T.F., Data Curation, S.I., T.U. and J.U.; Writing-Original Draft, S.I.; Writing-Review & Editing, T.F., T.U., and J.U., Visualization, S.I.; Supervision, S.I., and J.U., Project Administration, S.I., and J.U.; Funding Acquisition, S.I., and J.U.

