## Supplementary Information for "Enhanced human sensorimotor integration via self-modulation of the somatosensory activity"

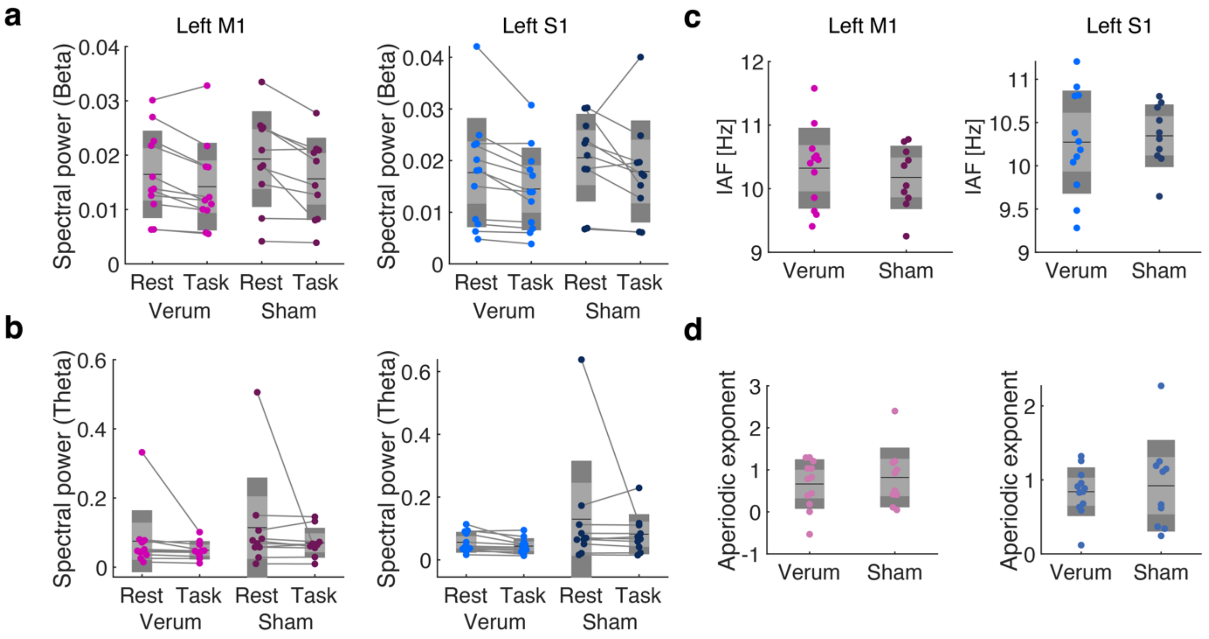


**Supplementary Figure 1 Analysis for power spectral density features.**

**a. Spectral power** at the beta-band at the rest and task periods in left M1 and S1. Each dot represents an individual participant. The mixed rmANOVA for M1 and S1 data indicated a significant main effect of condition (Rest and Task) (M1: $F = 22.0$, $p < 0.001$, $\eta^2 = 0.035$; S1: $F = 6.38$, $p = 0.02$, $\eta^2 = 0.026$). The post-hoc *t*-test revealed the significant decrease in the beta-band power at the task period (M1: $t = 4.69$, $p < 0.001$, $d = -0.37$; S1: t = 2.53, $p = 0.02$, $d = -0.32$). **b. Spectral power** at the theta-band at the rest and task periods in left M1 and S1. No significant main effects or interaction were found in both M1 and S1 data (M1: Condition, $F = 2.2$, $p = 0.15$, Group, $F = 1.1$, $p = 0.31$, Interaction: $F = 0.14$, $p = 0.71$; S1: Condition, $F = 1.55$, $p = 0.23$, Group, $F = 2.97$, $p = 0.1$, Interaction: $F = 0.53$, $p = 0.47$). **c.** Individual Alpha Frequency (IAF) comparison between Verum and Sham groups in left M1 and S1. **No evidence in the systematic difference between groups were found in the left M1 or S1** (M1: $t = -0.21$, $p = 0.83$; S1: $t = -0.36$, $p = 0.72$)**.** **d.** Aperiodic exponent comparison between Verum and Sham groups left M1 and S1. **No evidence in the systematic difference between groups were found in the left M1 or S1** (M1: $t = -0.42$, $p = 0.70$; S1: $t = -0.52$, $p = 0.61$)**.**

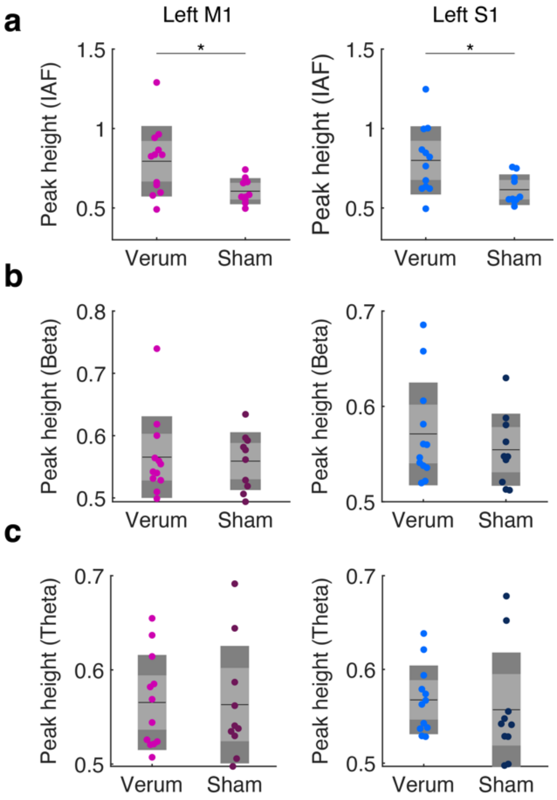


**Supplementary Figure 2 Analysis for the parameterized spectral peak height.**

**a.** Comparison of spectral peak heights at the individual alpha frequency (IAF) range for the Verum and Sham groups in the left M1 and S1. A significant difference in peak height is observed during the test period between the verum and sham groups in both regions (Two-sample $t$-test, Left M1: $t = 2.79$, $p = 0.0014$, $d = 1.15$; Left S1: $t = 2.74$, $p = 0.0015$, $d = 1.14$). The peak height was parameterized by the *specparam* algorithm. **b.** Comparison of spectral peak heights at the beta-band. (Two-sample $t$-test, Left M1: $t = 0.09$, $p = 0.93$; Left S1: $t = 0.49$, $p = 0.64$). **c.** Comparison of spectral peak heights at the theta-band. (Two-sample $t$-test, Left M1: $t = 0.27$, $p = 0.79$; Left S1: $t = 86$, $p = 0.40$).

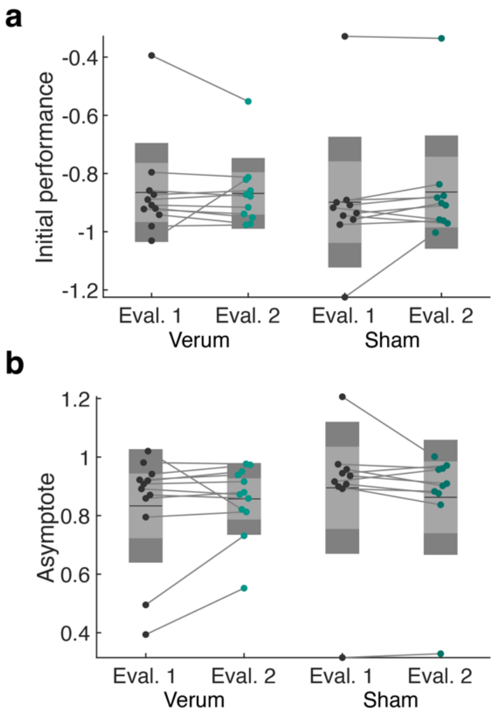


**Supplementary Figure 3 Analysis for motor performance parameters.**

**a.** Comparison of initial performance change for the Verum and Sham groups parameterized by the exponential fitting. The parameter *a* reflects the performance at the shorter movement time. No significant interaction and main effects were found by the rmANOVA analysis. **b.** Comparison of asymptote change for the Verum and Sham groups parameterized by the exponential fitting. The parameter *c* reflects the performance at the longer movement time. No significant interaction and main effects were found by the rmANOVA analysis.

*Script for the instruction of kinesthetic motor imagery*

First, perform abduction of the index finger. At this time, make sure that the muscles in your finger are engaged. Next, perform the same movement with about half the strength as before. Again, ensure that the muscles in your finger are engaged. Then, perform the same movement with about half the strength as in the previous step. Finally, without moving your finger, imagine the sensation you felt when performing the previous movements. You will repeat this mental imagery, referred to as kinesthetic motor imagery, during the task.Be careful not to actually engage the muscles in your finger during this process.
